# The Neuron Phenotype Ontology: A FAIR Approach to Proposing and Classifying Neuronal Types

**DOI:** 10.1101/2020.09.01.278879

**Authors:** Thomas H. Gillespie, Shreejoy Tripathy, Mohameth François Sy, Maryann E. Martone, Sean L. Hill

## Abstract

The challenge of defining and cataloging the building blocks of the brain requires a standardized approach to naming neurons and organizing knowledge about their properties. The US Brain Initiative Cell Census Network, Human Cell Atlas, Blue Brain Project, and others are generating vast amounts of data and characterizing large numbers of neurons throughout the nervous system. The neuroscientific literature contains many neuron names (e.g. parvalbumin-positive interneuron or layer 5 pyramidal cell) that are commonly used and generally accepted. However, it is often unclear how such common usage types relate to the many proposed evidence-based types that are based on the results of new techniques. Further, comparing different models across labs remains a significant challenge. Here, we propose an interoperable knowledge representation, the Neuron Phenotype Ontology (NPO) that provides a standardized and machine computable approach for naming and normalizing phenotypes in cell types by using community ontology identifiers as a common language. The NPO provides a framework for systematically organizing knowledge about cellular properties and enables interoperability with existing neuron naming schemes. We evaluate the NPO by populating a knowledge base with three independent cortical neuron classifications derived from published data sets that describe neurons according to molecular, morphological, electrophysiological and synaptic properties. Competency queries to this knowledge base demonstrate that this knowledge model enables interoperability between the three test cases and common usage neuron names from the literature.

## Introduction

The modern description and classification of neurons and the diversity of their properties began with the work of Santiago Ramon y Cajal over 100 years ago. Cajal benefitted from a newly discovered technique, the Golgi stain, to reveal neurons as individual entities of remarkably different shapes, which he described as the “ butterflies of the soul”. Our knowledge of neuron types (as with cell types) has continued to evolve as new experimental techniques emerge. For this reason, a centerpiece of the US Brain Initiative is to re-examine what constitutes a cell type in light of new ways of probing the nervous system. Through the BRAIN Initiative Cell Census Network (BICCN) researchers are generating large pools of data using cutting edge methods that are being integrated across data types through the use of standards such as a common spatial and semantic mappings (Ecker et al. 2017). The BICCN joins several other large initiatives such as the Blue Brain Project (Markram 2006), Human Cell Atlas (Regev et al. 2017), and SPARC (https://sparc.science/) which also seek to provide foundational knowledge on the types of cells that make up the nervous system. As these data are analyzed and synthesized, new ways to distinguish among different classes of neurons are being proposed and published.

One of the end goals of these large projects is to integrate and analyze large quantities of cellular data to derive new taxonomic classification of neurons across neural structures and to arrive at a new understanding of what constitutes a cell type in the nervous system. To manage this process, some have called for a consistent naming scheme for neurons, so that as new types are discovered, their findings can be reported and compared in an organized way (Hamilton et al., 2012 ; DeFelipe et al. 2013; Shepherd et al. 2019). Biology has a long history of successfully developing and deploying taxonomies and naming conventions for new entities, e.g., species, enzymes. The process usually involves the commissioning of an authoritative body that comes up with a regularized method and vocabulary for distinguishing among different types and applying an appropriate nomenclature. However, developing either of these pre-supposes that we understand the key dimensions across which neurons should be classified and the foundations of what constitute a cell type. For example, the Petilla terminology proposed a set of criteria and controlled terminology for naming cortical interneurons (Petilla Interneuron Nomenclature Group et al. 2008). In addition, as new technologies enable further characterization of additional dimensions, our concept of cell types is likely to evolve. On the other hand, although we expect new insights regarding defining cell types in the nervous system, to date large integrative data gathering exercises have tended to refine our current concepts rather than replace them (Osumi-Sutherland 2017). In a single cell transcriptomic analysis of retinal bipolar cells, (Shekhar et al. 2016), detected 17 different types of RBC, 15 of which corresponded to those previously described. The challenge remains to define a knowledge representation that can readily adapt to and integrate results from new data-driven taxonomic efforts but which still supports references to classical naming schemes to ensure integration with the large amount of historical published knowledge.

Most proposed schemes, to date, comprise a hierarchical method based on various phenotypic properties for their foundation, i.e., key molecular, physiological, and connectivity signatures that distinguish a neuron type. Phenotypic properties are typically properties of a neuron which are consistent across a variety of measurements, although many phenotypic properties can only be consistently reproduced with a specific experimental technique or protocol. Given the multiple dimensions across which neurons can be differentiated, a phenotype-based approach for classification could effectively generate an almost infinite number of ways to categorize neurons, depending on the granularity at which the distinctions are expressed. A single taxonomy that effectively organizes neurons across these dimensions is unlikely. The recent proposal for naming cortical neurons by (Shepherd et al. 2019) shows how quickly the number of phenotypes can explode, particularly when trying to address the results of dense phenotypic sampling such as array expression. Thus for neuronal cell types, given the complexity and variety of potentially distinguishing features and the likely evolution of these over time, any system for communicating and comparing across phenotypes will require a firm computational foundation.

Traditionally, such proposed classifications are communicated through the research paper, where any taxonomy proposed is presented in the form a table, dendrogram or some other figure (e.g., Paul et al., 2017, Table S7; Markram et al., 2015, Table 1). The problem with our traditional way of constructing and communicating these taxonomies is that they require a human being to understand, compare, and reconcile them (Petilla Interneuron Nomenclature Group et al. 2008). Anyone who has attempted to read through multiple articles, each with their own proposal for classifying cell types within a region understands the difficulties in trying to reconcile the different schemes, even when they are based on limited numbers of data dimensions. The multiplicity of papers proposing classification schemes just for cortical interneurons illustrates this point (Cauli et al. 1997). With the BICCN and other large scale consortia tasked to map the cellular landscape of the brain and body, the potential number of these taxonomies is likely to explode beyond the current already unmanageable number, as researchers apply new types of analytics to understand the data. For neuroscience to move beyond paper-based forums for discussion and integration, we need to treat taxonomies and names as computable artifacts that comply with the FAIR data principles, FAIR =Findable, Accessible, Interoperable and Reusable; (Wilkinson et al. 2016).

**Table 1.** Phenotypic Dimensions of the NPO and the associated ontologies/vocabularies used to populate the data model

| Phenotypic dimension | Definition | Vocabularies/ontologies |
| --- | --- | --- |
| Organism | The species or taxon rank in which the phenotype inheres | NCBI taxonomy<sup>1</sup> |
| Anatomical | The region of the nervous system containing parts of the neuron. Primary location is indicated by the location of the cell soma, but anatomical location may be assigned to any cell part through a series of predicates | UBERON; various brain atlases via NIFSTD parcellation<sup>2</sup> |

<sup>1</sup> https://www.ncbi.nlm.nih.gov/taxonomy
<sup>2</sup> https://github.com/SciCrunch/NIF-Ontology/blob/master/docs/brain-regions.org

|  |  |  |
| --- | --- | --- |
| Morphological | Distinguishing morphological characteristics | NIFSTD<sup>3</sup> |
| Molecular | Distinguishing molecular constituents | NCBI Gene<sup>4</sup>, CHEBI<sup>5</sup>, Protein Ontology<sup>6</sup> |
| Physiological | Expresses a relationship between a neuron type and an electrophysiological phenotype concept. This should be used when a neuron type is described using a high level electrophysiological concept class, e.g., bursting | Petilla Conventions (Petilla Interneuron Nomenclature Group, 2008) |
| Connection | Indicates a synaptic relationship between cell types. Further elaborated into connectivity determined by different techniques, e.g., physiology, electron microscopy | Gene Ontology<sup>7</sup> |
| Circuit role | Indicates whether the neuron is an Intrinsic neuron (local circuit neuron), projection neuron or sensory neuron | NIFSTD (Bug et al., 2008) |
| Projection targets | Expresses a relationship between a neuron type and a brain region to which it sends axons. Synaptic relationships are represented through the connection relationship. | UBERON (Mungall et al., 2012)/various atlases/NIF Gross Anatomy (Bug et al., 2008) |

Towards that end, we have developed an ontology-based data model, the Neuron Phenotype Ontology (NPO). The NPO aims to provide an interoperable representation of cell types that can evolve as our phenotypic knowledge evolves, from initial data gathering to modeling and synthesis (Fig 1). The NPO provides a computable representation of cell types defined by collections of phenotypic properties, designed to enable interoperability between neuronal taxonomies. It is designed to enable scientists to discover which cell types (or potential cell types) share similar properties and to help scientists understand when the cell types they observe are the same or similar to other cell types described in the literature or from other laboratories. Here, we show how the NPO can be used to express taxonomies proposed by different research groups using modern techniques, enable comparisons between them, and enable queries with commonly used neuron types from the literature.

**Figure 1.**
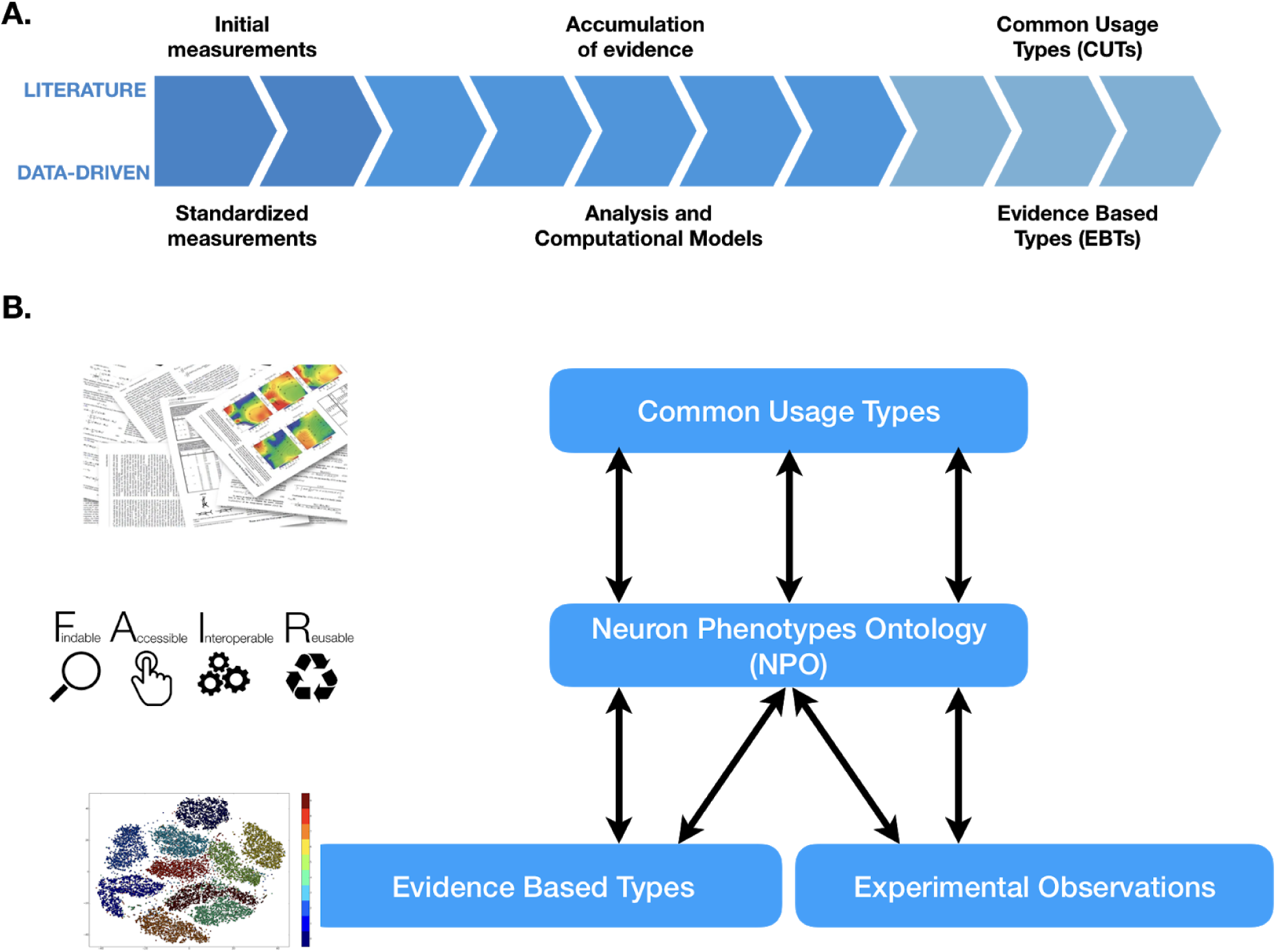
Evolution of neuron knowledge. A. Common usage types (CUTs) emerge in the literature as evidence accumulated for generally accepted neuron types with implicitly known properties. Data-driven studies generate evidence-based types (EBTs) based on explicitly measured standardized properties. B. The Neuron Phenotype Ontology (NPO) provides interoperability between the CUTs from the literature, the EBTs from data-driven studies, and new experimental observations from individual laboratories.

## Methods

### Overview of NPO

The NPO as well as all data and code referenced below are available for reuse under open licenses (see Data and Code availability statement).

The NPO provides a data model for modeling a neuron type as a “ bag of key phenotypes”, that is, neurons are represented as a collection of phenotypic properties (Fig 2) formalized by Web Ontology Language (OWL) classes. These properties can then be used to communicate about and compare phenotypes across laboratories, species and experimental techniques. This approach has been demonstrated previously in the context of text-based queries of neuron type mentions (Richardet et al. 2015).

**Figure 2.**
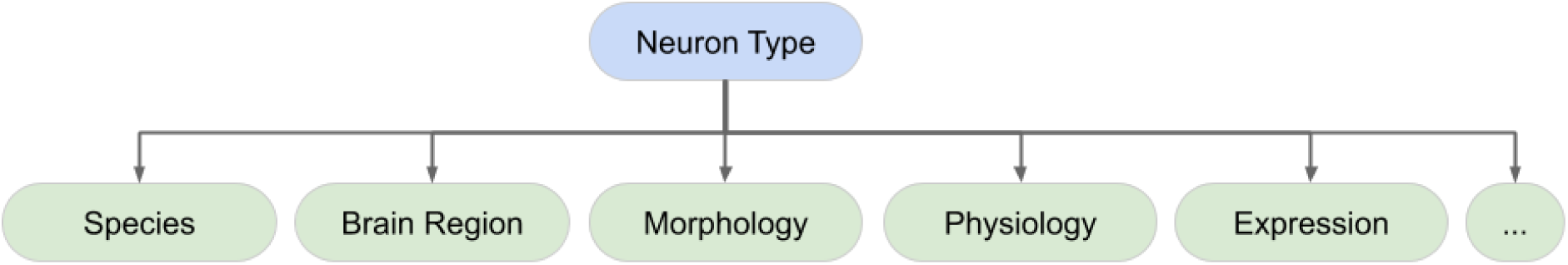
High level data model for neuron phenotypes. The Neuron Phenotype Ontology characterizes neuron types as bundles of normalized phenotypic properties.

Each of these dimensions is linked to a formal vocabulary or ontology, which is used to provide the descriptors for qualitative phenotypic attributes (Table 1). When possible, the vocabularies are drawn from community ontologies/vocabularies in broad use across biomedicine to aid in interoperability. Those dimensions that were not covered by specific community ontologies were added as classes to the appropriate branches of the NIFSTD ontology. NIFSTD is a harmonized set of neuroscience relevant ontologies developed and maintained by the Neuroscience Information Framework (Bug et al. 2008). These dimensions are further elaborated in a set of predicates that capture more granular aspects of phenotypes. For example, *hasMolecularPhenotype* can be further divided into *hasNeurotransmitterPhenotype, hasEpigeneticPhenotype*, and *hasExpressionPhenotype* (Fig 3). *hasExpressionPhenotype* is further broken down into a set of predicates that captures the methodology used to reveal the phenotype. In the current version (v1) of the NPO, we have not made use of the full set of relationships to simplify the reasoning. Relationships that have not been used in the current version of the NPO are grayed out in Figure 3.

**Figure 3.**
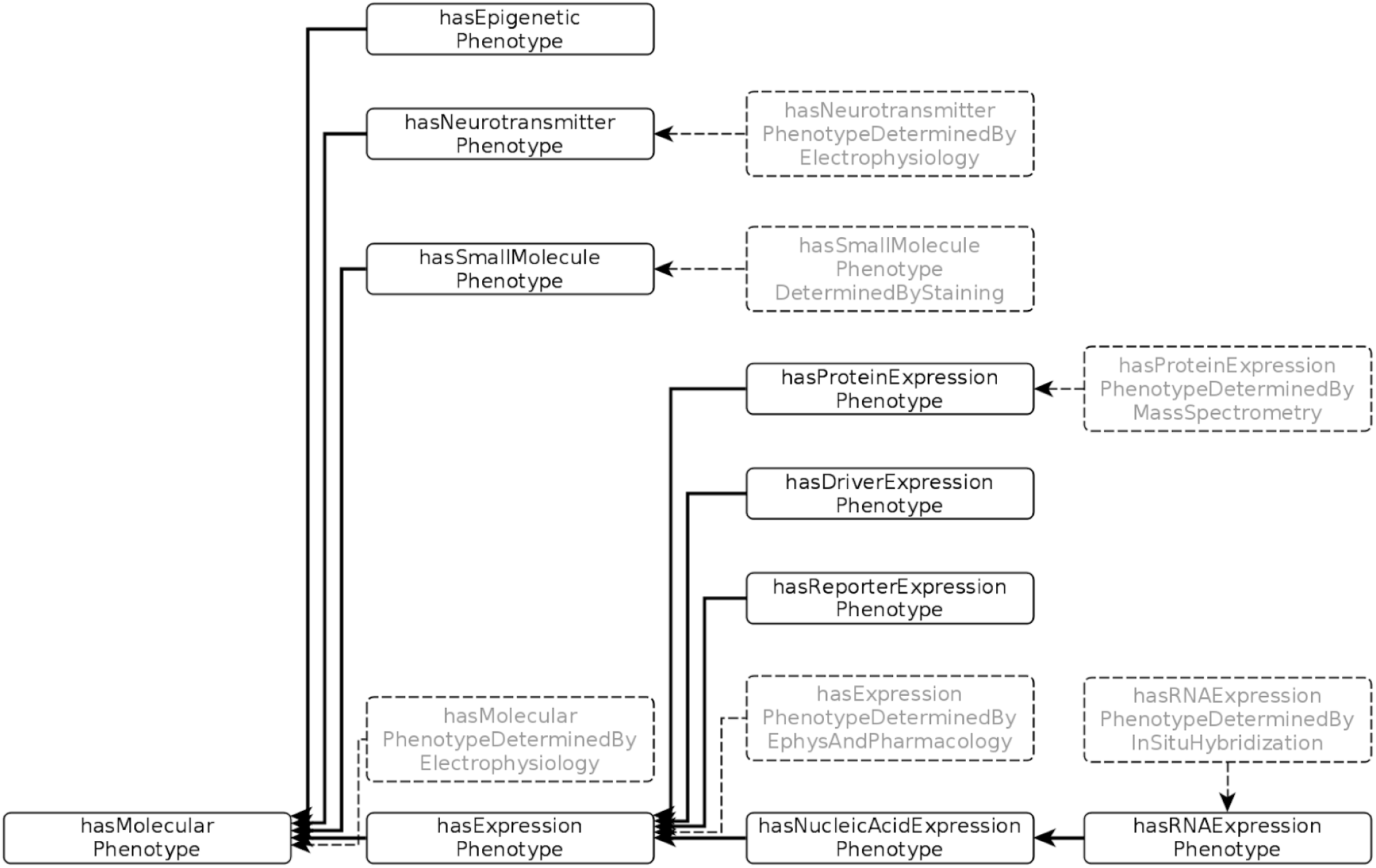
The set of predicates employed to define molecular phenotypes

For negative phenotypes, that is, where the lack of a particular phenotype is considered to be a distinguishing feature between neuron types, we use negation in OWL semantics, e.g., a parvalbumin negative neuron would be modeled as “ not (*hasExpressionPhenotype* some ‘parvalbumin alpha’)”.

We have also included disjointness axioms in cases where the strength of the assertions from the EBTs were not as definitive as full negation.

For evaluation purposes, we have used the NPO data model to construct a knowledge base of neuronal phenotypes comprising two branches: 1. Phenotypic representations of common usage types (CUTs) from classical morphological and physiological studies over the past 100 years; 2. Classification models arising from newer experimental techniques tied to individual projects,laboratories or initiatives, termed evidence based types (EBTs). The data model is supported by computational tools that enable individual researchers to compose the complex phenotype of a neuron out of any number of individual phenotypes that are tightly linked to individual data sets and analyses (Fig 4). Neuron Data Model (NeuroDM) is a python library that implements Neuron Lang, a domain specific language (DSL) for specifying neurons and generating human readable neuron names based on these OWL semantics. NeuronDM provides tools for mapping to and from collections of local names for phenotypes by using ontology identifiers as the common language underlying all local naming. These tools also let us automatically generate names for neurons in a regular and consistent way using a set of rules operating on the neurons’ constituent phenotypes. Neuron Lang can export to python or to any serialization supported by rdflib, however deterministic turtle^8^ (ttl) is preferred.

**Figure 4:**
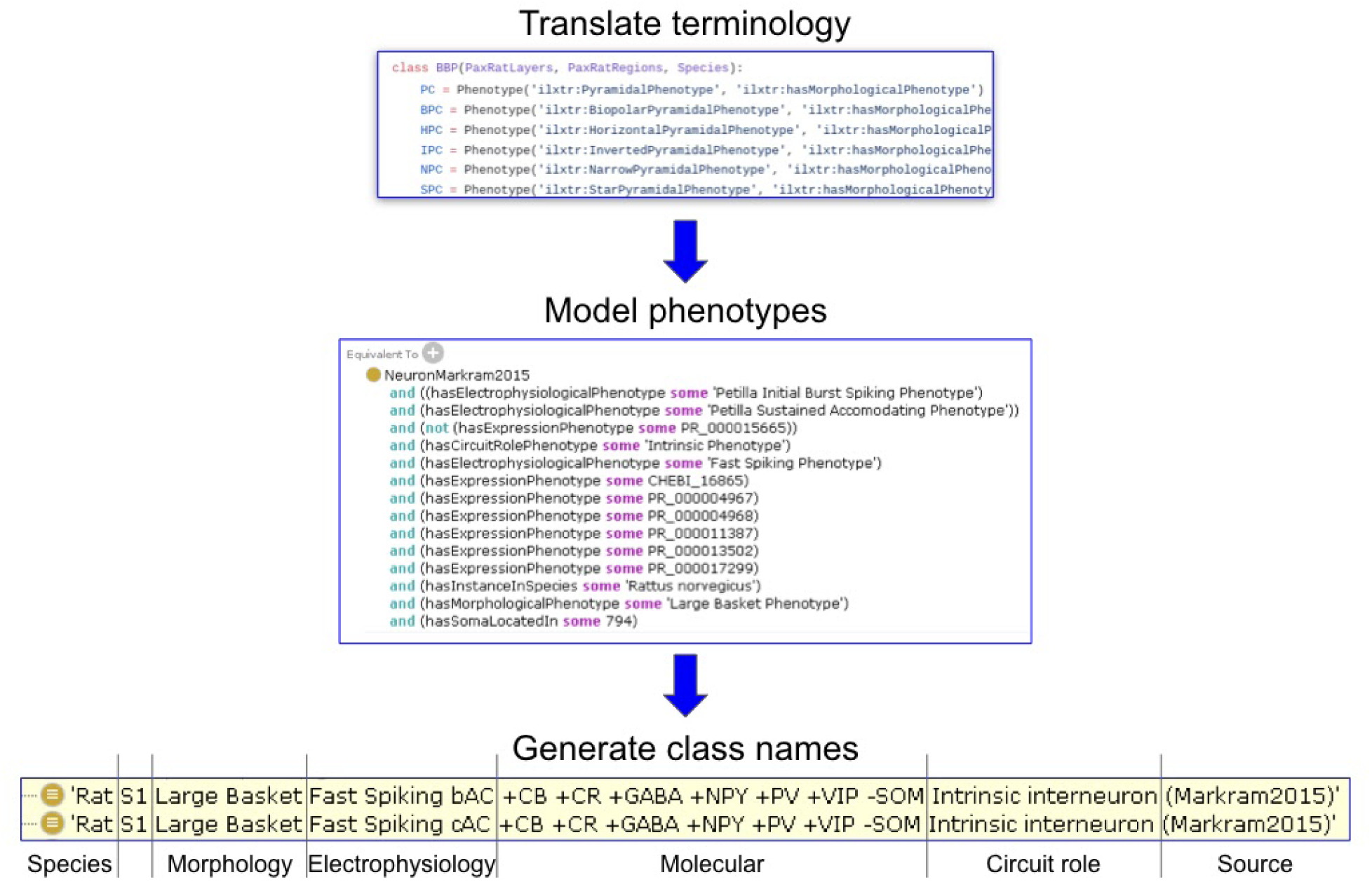
Process used to translate local terminology into ontology-based representations and machine-generated names. Using NeuronDM, phenotypes are first mapped by a user into ontology identifiers (top panel). NeuroDM automatically translates these mappings into OWL equivalence statements (middle panel). Neuro Lang then generates a set of human readable labels based on these restrictions (bottom panel).

### Modeling decisions

#### Neuron class names

Each neuron in the NPO is identified by a full uniform resource identifier (URI) and a compact identifier for ease of reference. The compact identifier has the prefix npokb and the ontology is registered in BioPortal^9^ (RRID:SCR_002713) using the NPOKB prefix as NPO prefix was taken. Each class has multiple human readable labels assigned as annotation properties. Neurons are named according to the phenotypic properties they display. These labels are generated automatically based on the collection of phenotypic properties reported for each cell type using the neuronDM Python library. Phenotypes are expressed as OWL2.0 restrictions, and neuron types as equivalent to the intersection of those restrictions (Fig 4). NPO provides two versions of these names. *Local label* records molecular properties in the native form in which they were measured, e.g., genes, proteins, transgenes, while the *rdfs:label* contains a normalized view where molecules are assigned a common molecular abbreviation regardless of the form in which it was measured (see below). For ease of reference we also preserve the common name for the CUT and the original name assigned by the investigator for EBTs if it was provided. These can be found under *origLabel*, and they also appear as *skos:prefLabel* when they are present, otherwise *skos:prefLabel* is populated from *rdfs:label* so there are no neurons missing a preferred label.

For the NPOKB, we generally follow the ordering recommended by (Hamilton et al. 2012) and (Ecker et al. 2017). In both papers, the recommendation was to create an ordered taxonomy based on key phenotypic features, arranged roughly hierarchically, starting from the highest level, species, followed by anatomical regions, then a set of standardized names for morphological, physiological, molecular or connectional phenotypes (Fig 4 lower panel). In this way, as proposed originally by Hamilton et al., (2012), it is easy to generate a human readable list of neurons from a given species or brain region and to compare across complex phenotypes. In addition, while we are still sorting out what constitutes a cell type, we define the local environment in which the neuron resides.

#### Molecular Indicators

For EBTs, NPO preserves the means by which molecular phenotypes are determined. If gene expression is measured, we use the identifier for the gene; if the expression of a transgene is measured, we include the transgene; if the protein is measured, we include the protein. For CUTs, we only use the protein, peptide or small molecules that are thought to define the class. In order to tie together these different measurements, we created a class called phenotype indicator (PhenotypeIndicator) that groups together the different forms of molecular entities, e.g., a somatostatin indicator is equivalent to Sst, SST, SOM, Sst-IRES-Cre, Sst-IRES-FLpO. A somatostatin neuron is then defined as equivalent to any neuron that has some somatostatin indicator as a molecular phenotype. In this way, we simplify the reasoning required to retrieve all somatostatin neurons, but we also clearly preserve the statements made by investigators in their instances or model assertions as preserved in the l*ocalLabel*. In addition, to translate all of the different representations of a particular molecular entity into a consistent human readable label, we have assembled a set of short names that represent each class based largely on common conventions or the names used in NCBI for mouse genes. These short names are used in the skos:hiddenLabel for each class and are suffixed with “ (indicator)” to create the *rdfs:label*. For example, when generating a label phenotype indicators for parvalbumin are shown as PV. These labels are available through the “ hidden label” annotation property under the ilxtr:PhenotypeIndicator class.

## Data and Code Availability

The NPO/NPOKB can be viewed by loading the .ttl file available at https://raw.githubusercontent.com/SciCrunch/NIF-Ontology/neurons/ttl/npo.ttl into the Protégé Ontology Tool (RRID:SCR_003299) v5.5.0 or higher. As described in the supplemental methods, the .ttl file is “ light” version of the full ontology that makes it less reliant on the full import chain. Additional information about working with the NPOKB can be found in the supplemental methods. The NPO is distributed under a CC-BY 4.0 Attribution license, but it imports community ontologies that may be covered under different licenses.

The work here describes v1.0 of the NPO which can be accessed at https://raw.githubusercontent.com/SciCrunch/NIF-Ontology/npo-1.0/ttl/npo.ttl. In the import closure of npo.ttl there are no external imports except for http://purl.obolibrary.org/obo/bfo.owlwhich had version Iri http://purl.obolibrary.org/obo/bfo/2019-08-26/bfo.owl at the time npo 1.0 was released. All other ontology iris resolve to the neurons branch of the NIF-Ontology except for http://ontology.neuinfo.org/NIF/ttl/generated/parcellation-artifacts.ttl. As a result, importing npo.ttl directly in Protégé will result in the newest version of the imports on the neurons branch being used, which may lead to some small differences in the results compared to what are presented here.. However, it is possible to use the NIF-Ontology catalog file to load an exact view of version 1.0 of npo.ttl by cloning the git repository and checking out the npo-1.0 tag.

The NPOKB is available on BioPortal at https://bioportal.bioontology.org/ontologies/NPOKB. A loaded graph that can be used with SciGraph, a neo4J-based database for serving ontologies, is available at https://github.com/SciCrunch/NIF-Ontology/releases/tag/npo-1.0.

The content of the NPO is also accessible via the UCSD SciCrunch SciGraph API at https://scicrunch.org/api/1/sparc-scigraph/. Documentation for access can be found at http://ontology.neuinfo.org/docs/NIF-Ontology/README.html#using-nifstd.

The neurondm git repo is https://github.com/tgbugs/pyontutils/tree/master/neurondm.

All python code bears an MIT license and is available on pypi. It can be installed via ‘pip install neurondm’. Additional instructions are available in the README.

Gentoo linux ebuilds for installing neurondm are available in the tgbugs-overlay. It can be installed via ‘layman -a tgbugs-overlay && eselect repository tgbugs-overlay && emerge neurondm’.

An archive of the code corresponding to this publication is also available on Zenodo at https://doi.org/10.5281/zenodo.4005727. Additional release artifacts are also available on the GitHub release page https://github.com/tgbugs/pyontutils/releases/tag/neurondm-0.1.3.

The full list of CUTs is available at: https://github.com/tgbugs/pyontutils/releases/download/neurondm-0.1.3/data-bundle-2020-08-28.zip

The full datasets produced for the competency queries (see Results) are available at: Gillespie, Martone, and Hill (2020) https://zenodo.org/record/4007065#.X03TD2dKiAZ

## Results

### Common Usage Types

Common usage types represent neuron types that have been reliably identified over many years by multiple groups using multiple techniques. The criteria we used to identity CUTs is provided in Supplementary Table S1. A master spreadsheet was created in Google Spreadsheets and populated with a list of neuron “ stubs” that were created automatically by taking the list of major brain regions in the UBERON ontology and creating two classes per region: Region X projection neuron and Region X intrinsic neuron. These anatomical regions were at a fairly coarse level and comprised the major brain and spinal cord regions, but generally not subregions, for example, cerebral cortex and not motor cortex. Individual brain regions were then augmented with the list of neuron types extracted from online knowledge bases. We started with the list of approximately 300 mammalian neurons from Neurolex Wiki (RRID:SCR_005402; (Larson and Martone 2013) that had been compiled through expert input via the Neuron Registry Task Force of the INCF (Hamilton et al. 2012), as well as by community contributions. This list was then cross referenced to NeuroElectro (RRID:SCR_006274), BAMS Cells (RRID:SCR_003531), Hippocampome.org (RRID:SCR_009023), NeuroMorpho.org (RRID:SCR_002145) and Blue Brain Project (RRID:SCR_002994). All of these sources were accessed via the Neuroscience Information Framework (RRID:SCR_002894) project to find a set of cells that were referenced in multiple databases. As NeuroElectro maps their nomenclature to the Neurolex names, we used this database to examine representation of these cell types in the neurophysiology literature. We selected all neurons that were referenced in more than one paper.

This procedure resulted in a working list of ∼350 neurons (for full list see Data Availability Statement). From this list, we then selected ∼100 neurons for which we had basic morphological and molecular properties available. We also included the neurotransmitter for the majority. We elected to focus in v1.0 primarily on molecular and morphological phenotypes, rather than the full complexity available in the NPO (Fig 2), as these are the most well known for CUTs and are the most frequent types encountered in the EBTs (Zeng and Sanes, 2018). We also elected in the modeling to take a minimalist approach, that is, our representation is meant not to represent an exhaustive list of every molecule that has been identified within a neuron, but the minimum set of molecules and morphological features that are characteristic for that type. This decision allowed us to construct OWL equivalence statements for each CUT that defined the necessary and sufficient conditions that would allow EBTs to classify under these CUTs. Additional phenotypes were still recorded but added through the Subclassof axiom. Subclassof represents a weaker form of restriction, representing a necessary but not sufficient condition for membership in a class. In order to avoid logical inconsistencies that would interfere with classification, we only included positive phenotypes in necessary and sufficient conditions for CUTs. If distinguishing negative phenotypes were present, they were modeled as entailments rather than OWL restrictions.

Following (Larson et al. 2007), the primary anatomical location of a neuron is assigned based on the brain region in which the soma is located, e.g., cerebellar neuron is equivalent to a neuron with a cell soma in any part of the cerebellum.

### Evidence-based types

EBTs represent cell types and taxonomies proposed by a single group based on an analysis of experimental evidence. For this version of the NPO and for the purposes of evaluating our phenotype model, we focused on 3 projects that have generated cortical classifications based on large amounts of experimental data:

A. **Cortical cell types proposed by the Blue Brain Project (Markram et al. 2015), as elaborated in the text and Table 1**. In this study, 56 total types across 9 morphological types are identified and physiologically characterized from cells in cortical area S1 of rats ranging from P11-P15 from which they recorded physiological properties. Cell-specific molecular markers were confirmed by immunohistochemistry and RT-PCR. (Markram et al. 2015) utilize a nomenclature aligned to the Petilla conventions (Petilla Interneuron Nomenclature Group et al. 2008) to annotate their physiological properties. For NPO V1.0, we included the molecular, morphological and electrophysiological phenotypic dimensions.
B. **The classification of proposed cortical GABAergic cell types from Josh Huang and colleagues as summarized in Table S7 of Paul et al**. (**2017**) **supplemented with additional information from Fig 1**. The latter was used primarily to create disjointness axioms (see Fig 1b). For NPO v1.0, we concentrated primarily on the gene expression phenotypes presented in this table, supplemented with information from the rest of the paper, e.g., disjointness axioms based on Fig 1b. Synaptic and physiological phenotypes will be included in a later version.
C. **The ∼800 cell classes contained in the Allen Cell Types database (RRID:SCR_014806), a database of experimental electrophysiological, morphological and transcriptomic data derived from single cell data**. In the Cell Types database, no classification scheme was proposed; rather the records represent statistical summaries of properties measured from these classes of cells identified in transgenic lines. We therefore include this as an EBT. For this version, we focused on molecular measurements from mouse cortex.

### Competency Queries

The NPO was designed to classify neurons according to phenotype dimensions, regardless of whether they represent EBTs or CUTs. To test the integrity of the knowledge base and the structure of the ontology, we developed a set of competency queries (CQ):

1. ***Find all parvalbumin+ neurons*** Description Logic (DL) Query: hasPhenotype some ‘parvalbumin (indicator)’
2. ***Find all cortical neurons that contain somatostatin*** DL Query: hasPhenotype some ‘somatostatin (indicator)’ and hasSomaLocatedIn some (neocortex or ‘part of’ some neocortex)
3. ***How do basket cells described in Paul et al. (2017) and Markram et al. (2015) compare on key dimensions?*** DL Query: (NeuronHuang2017 or NeuronMarkram2015) and hasPhenotype some ‘Basket phenotype’
4. ***What EBTs are related to the Martinotti cell?*** Determine which neurons classify under the CUT Neocortex Martinotti cell

All of the results presented below were produced by issuing OWL DL queries as specified above in Protégé v5.5.0 on a MacBook Pro using the ELK 0.4.3 reasoner unless otherwise noted. More information on loading the ontology into Protégé can be found in the Supplemental Methods.

#### CQ1: Find all examples of parvalbumin neurons

This query should return all neurons that have a phenotype associated with parvalbumin, regardless of exactly what molecule was measured (DNA, RNA, protein) or how it was measured. In this version of the NPO, we achieve this by creating phenotype indicators without specifying the relationships between these measures through the npokb:parvalbumin (indicator) class. The results of this query are summarized in Table 2. A total of 86 neurons are returned, including EBTs (Huang, N = 2, Markram, N = 16 and Allen; N =59) and CUTs (N = 9). To aid in comparison across these classes, we illustrate with one example each from the Markram EBTs and Allen data. The complete list of neurons is provided in Gillespie et al., (2020). The original label is provided for each EBT and the common name for the CUT. These are followed by the *localLabel* names that preserve the form of molecule upon which the classifications were based to illustrate how the NPO can be used to compare across different assertions about molecular identity (Markram2015, Huang2017, AllenCT). Related phenotypic values are color coded to aid in comparison. In this case, we use the *localLabel* that preserves the form of molecule upon which the classifications were based. For a complete list of abbreviations, see Table S2.

**Table 2.** Examples of EBT and CUT neurons returned from Competency query CQ1: Find all examples of parvalbumin containing neurons. The form of the parvalbumin indicator is highlighted in red. Only one example is provided from the Allen EBT (total 59). Full results are available in Gillespie et al., 2020. The compact identifier for each class is prefixed (in bold) to the localLabel for ease of reference. The local label preserves the form in which the molecule was measured. The Common/original name represents the common name from the superclass for all of the physiological subtypes for the Markram cells. However, for the local label we provide a subtype as the superclass does not include the full molecular profile in the name.

| Type | # | Common/original name | NPO localLabel |
| --- | --- | --- | --- |
| CUT | 6 | nifext:56: Neocortex basket cell | Mammalia neocortex L2/3 Basket +PV<br>+GABA intrinsic neuron |
| **EBT Markram** | 16 | **npokb:112:**Nest basket cell | Rattus norvegicus S1 Nest basket (intersectionOf AC b) Fast spiking +GABA +calbindin +CR +NPY **+PV** +VIP -SST intrinsic neuron (Markram2015) |
| **EBT Huang** | 2 | **npokb:43:**PVBC cortical neuron | "Mus musculus neocortex Basket +GABA **+PV-cre** intrinsic neuron (Huang2017)" |
| **EBT Allen** | 59 | none | **npokb:434:**Mus musculus female left cerebral hemisphere VISrl2_3 -Apical Dendrite -Spiny **+Pvalb-T2A-FlpO** +Vipr2-IRES2-Cre +Ai65(RCFL-tdT) neuron (AllenCT) |

Three of the neuron classes indicate that the parvalbumin cells are basket cells, while the Allen data does not specify morphology beyond noting that these cells lack an apical dendrite and dendritic spines.

#### CQ2: Find all cortical neurons that contain somatostatin

This query should return all cortical neurons that contain somatostatin regardless of cortical subregion or atlas brain region. Details about how atlas brain regions are handled are provided in the supplemental methods. This query returns a total of 100 neurons, including the neocortex Martinotti cell from the CUT and EBTs from the three classification schemes (Table 4). For Markram, we show only one subtype from each of the 3 main types. For Allen, we selected a few representative examples. Note that Allen neurons are returned from retrosplenial cortex (RSPd2/3) and two areas of primary visual cortex (VISal6a, VISl5) while Markram is returned for primary somatosensory cortex (S1). Both Huang and Allen cells use a transgenic line for Sst expression, labeled SST and Sst-IRES-FlpO respectively. The local labels preserve the nomenclature used in the source (Paul et al 2017 and Allen Cell Types Database respectively). However, because the NPO maps to identifier systems wherever possible, we can see that Huang and Allen use the same transgenic line developed by the Huang lab, regardless of the different nomenclature (jax:028579). For the case of transgenes in NPO, the identifier is the Jackson lab stock number when it is available.

**Table 4.**
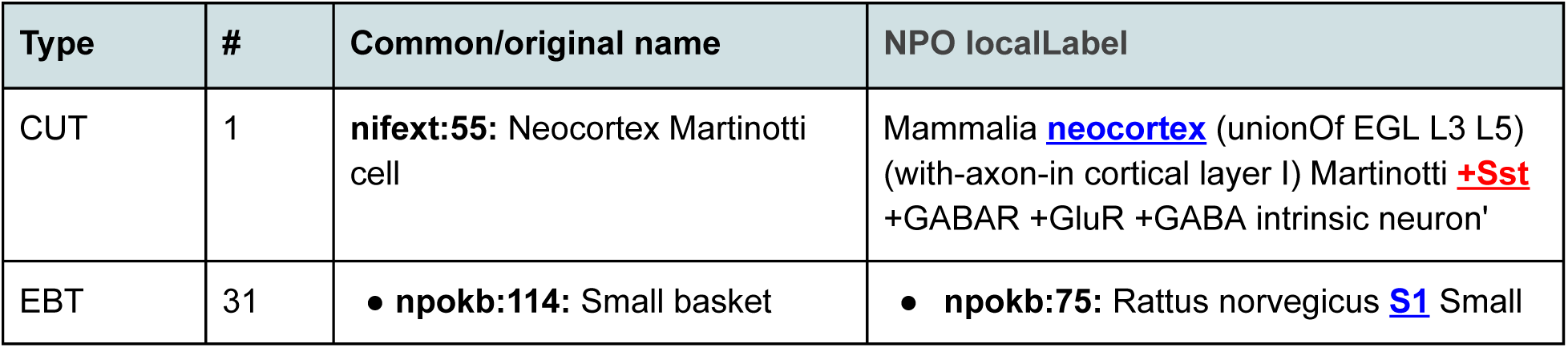
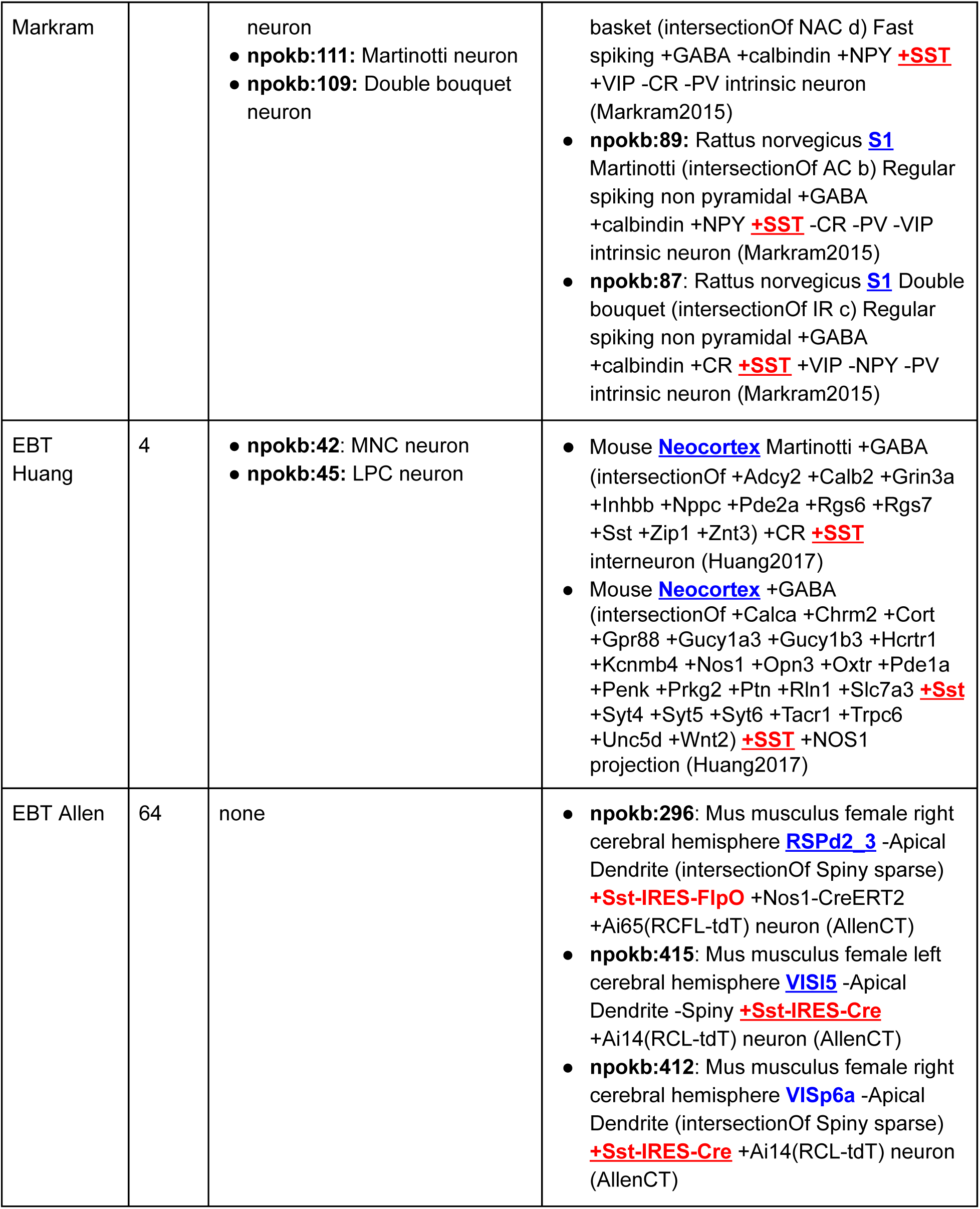
Results for CQ2: Find all cortical neurons containing somatostatin. Full results are available in Gillespie et al., (2020). The compact identifier for each class is prefixed (in bold) to the local label for ease of reference. The local label preserves the form in which the molecule was measured. The Common/original name represents the common name from the superclass for all of the physiological subtypes for the Markram cells. However, for the local label we provide a subtype as the superclass does not include the full molecular profile in the name. Similar entities across cell types are color coded. Brain region = blue; somatostatin indicator = red.

#### CQ3:How do basket cells described in Paul et al. (2017) and Markram et al. (2015) compare on key dimensions?

This query returned EBT cells from the two groups that were assigned the morphological phenotype “ basket”. A total of 22 neurons were returned, 20 from Markram and two from Huang. A subset are illustrated in Table 5 and related phenotypes are color coded across the different types for ease of comparison. For the Markram cells, we only show one subtype for each main class.

**Table 5.** Neurons that have a basket phenotype. Similar entities across the cell are color coded to aid in comparison. The full results list is available in Gillespie et al, 2020. Similar entities are color coded across cell types: blue = brain region; green = morphology; purple = neurotransmitter; dark red = parvalbumin indicator; red = somatostatin indicator.

| Original name | NPO ID | NPO Label |
| --- | --- | --- |
| PVBC Neuron (Huang2017) | npokb:43 | Mus musculus **neocortex** **Basket** **+GABA** (intersectionOf +Adm +Cckbr +PV +ilxtr:Kv3 +Rspo2 +Adcy8 +Cox6c +Gabra1 +Gabra4 +Gabrd +Gria1 +Gria4 +Mef2c +Pparg +Ppargc1a +Rgs4 +Slit2 +Slit3 +Tac1 +Arhgef10 +Esrrg +Nefh +Adcy1 +Rasl11b) **+PV** intrinsic neuron (Huang2017) |
| CCKC Neuron (Huang2017) | npokb:40 | Mus musculus **neocortex** **Basket** **+GABA** (intersectionOf +Crh +Cck +Cck +Cnr1 +Edn3 +Htr3a +Igf1 +VIP +VIP +Vipr1 +Adcy9 +Chrm3 +Cplx2 +Htr2c +Pnoc +Npy1r +Tac2 +Cplx3 +Pde7b +Prok2 +Hs6st3 +Syt10 +Rgs12) +Cck **+VIP** intrinsic neuron (Huang2017) |
| Large basket cell (Markram2015): subtype | npokb:59 | 'Rattus norvegicus **S1** **Large Basket** (intersectionOf AC b) Fast Spiking **+GABA** +Calb +Calb2 +Npy **+PV +VIP** -Sst interneuron (Markram2015)' |
| Nest basket cell (Markram2015): subtype | npokb:65 | 'Rattus norvegicus **S1** **Nest Basket** (intersectionOf AC b) Fast Spiking **+GABA** +Calb +Calb2 +Npy **+PV +VIP** -Sst interneuron (Markram2015)' |
| Small basket cell (Markram2015): subtype | npokb:73 | 'Rattus norvegicus **S1** **Small Basket** (intersectionOf AC c) Fast Spiking **+GABA** +Calb +Npy +Sst **+VIP** -Calb2 **-PV** interneuron (Markram2015)' |

Two classes of basket neurons are returned for Huang, while three are returned for Markram. Each of the three Markram classes are distinguished by distinct basket morphologies: small basket phenotype, large basket phenotype, and nest basket phenotype. These morphologies are modeled as subtypes of npokb:BasketPhenotype.

For these types of comparisons, the NPO facilitates comparison across diverse experimental techniques and anatomical nomenclatures and can help to generate testable hypotheses regarding phenotypes. In this example, it is difficult to tell from the information provided whether there is a 1:1 correspondence between any of the Huang and Markram cells. The only molecules mentioned by all 5 cells are GABA, PV and VIP. The Huang PVBC neuron is PV+ while the CCKC neuron is VIP+. Two Markram neurons are positive for both PV and VIP, while the small basket cell is asserted to be PV+ and VIP-. No negative phenotypes were recorded for the Huang neurons, as we based the equivalence classes on the information available in Table S7 which only included positive phenotypes. In the NPO, we operate under an open world assumption, that is, unless there is an explicit statement that a molecule is lacking, we do not assume that it is absent. We do provide additional information in the form of disjointness axioms based on Fig 1b of Paul et al. (2017) that the PV-containing and the VIP-containing cells are non-overlapping. This approach dovetails with EBTs making assertions about disjointness of cell types within a species which can be true even if there is not a universal axiom about molecular constituents. Disjointness therefore doesn’t mean that there is no expression, but an inspection of the data provided in Fig 1e indicates that expression of PV in the CCKC neuron is very low. Inspecting the data therefore suggests that the CCKC neuron is VIP+ and PV-, consistent with the small basket cell of Markram.

This example illustrates some of the difficulties involved in comparing across phenotypes, particularly when the different phenotypes are measured across experiments. It also illustrates the importance of tying EBTs to experimental data, so that predictions generated from these comparisons can be explored. In this case, Paul et al. (2017) provided expression data for several key molecules in FIg 1e. This figure shows that while the CCKC neuron expresses little to no PV, consistent with the small basket cell, it also expresses little to no Sst and detectable Calb2, in contrast to the small basket cell. However, as is easily seen in the labels, the Huang and Markram cells come from mouse and rat respectively and how complex molecular phenotypes compare across species is unknown (Yuste et al., 2020).

#### CQ4: What EBTs are related to the Martinotti cell?

To address this competency query, we reasoned over the ontology to determine which neurons would classify under the Neocortex Martinotti neuron CUT. For a neuron to be classified as a type of Martinotti cell, it has to share necessary and sufficient conditions of that class as coded in the equivalence statements. As discussed in the methods, we deliberately chose to model a minimum of properties as necessary and sufficient due to the large variability in the number of phenotypes recorded for the EBTs. Additional properties are included (Figure 5C) but not in the form of OWL restrictions, so they do not factor into the reasoning. We also only represent the major classes of CUTs and do not include subtypes, as these are less well agreed upon. In OWL, if we were to require that a Martinotti neuron must have calretinin, if a given EBT did not state that calretinin was a defining characteristic, the neurons would not classify. In fact, according to Rudy et al. (2011), Martinotti cells contain two subclasses, one that contains calretinin and one that does not. In the NPO, the NeuronHuang2017 EBT notes the presence of calretinin (+Calb2), while the NeuronMarkram2015 EBT says it is absent (-Calb2), perhaps representing these two subclasses.

**Fig 5.**
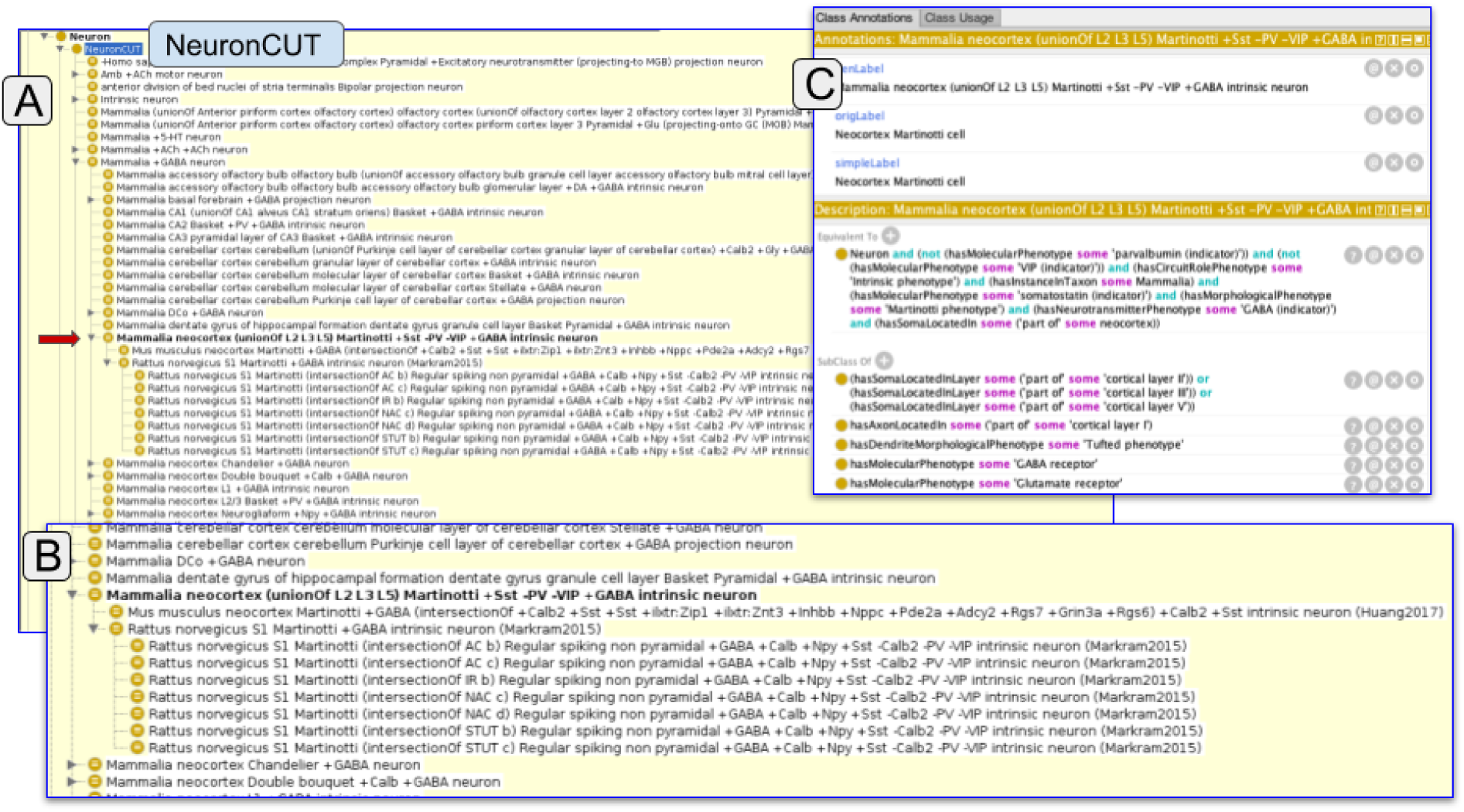
Inferred hierarchy after reasoning over the ontology for the Martinotti cell. Panel A shows the hierarchy generated under the NeuronCUT class. The position of the Marinotti CUT is indicated by the lower red arrow. An enlargement of the Martinotti classification is shown in panel B. Panel C shows the OWL representation of the Martinotti CUT.

As Fig 5 shows, the Allen EBTs do not classify under the Martinotti CUT. In v1.0 of the NPO, we only model morphological phenotypes at a coarse level, e.g., Martinotti phenotype, which is assigned to the level of the entire cell. In contrast, NeuronACT provided morphological information only for the dendrites of each cell. For the cortical somatostatin containing cells, it was noted that they lack an apical dendrite and dendritic spines, but no assertion was made about a Martinotti phenotype, unlike in the other two classifications. In the future, the NPO will include additional defining features of a Martinotti phenotype.

### FAIR properties of the NPO

The NPO was designed to be consistent with the FAIR principles. In Table 7, we show how the NPO achieves FAIR using the rubric in Hodson et al. (2018). The key features are machine readability, the use of identifiers (FAIR vocabularies), common knowledge representation languages and community standards. We provide a comparison with other cellular ontologies in Table S1.

**Table 7.**
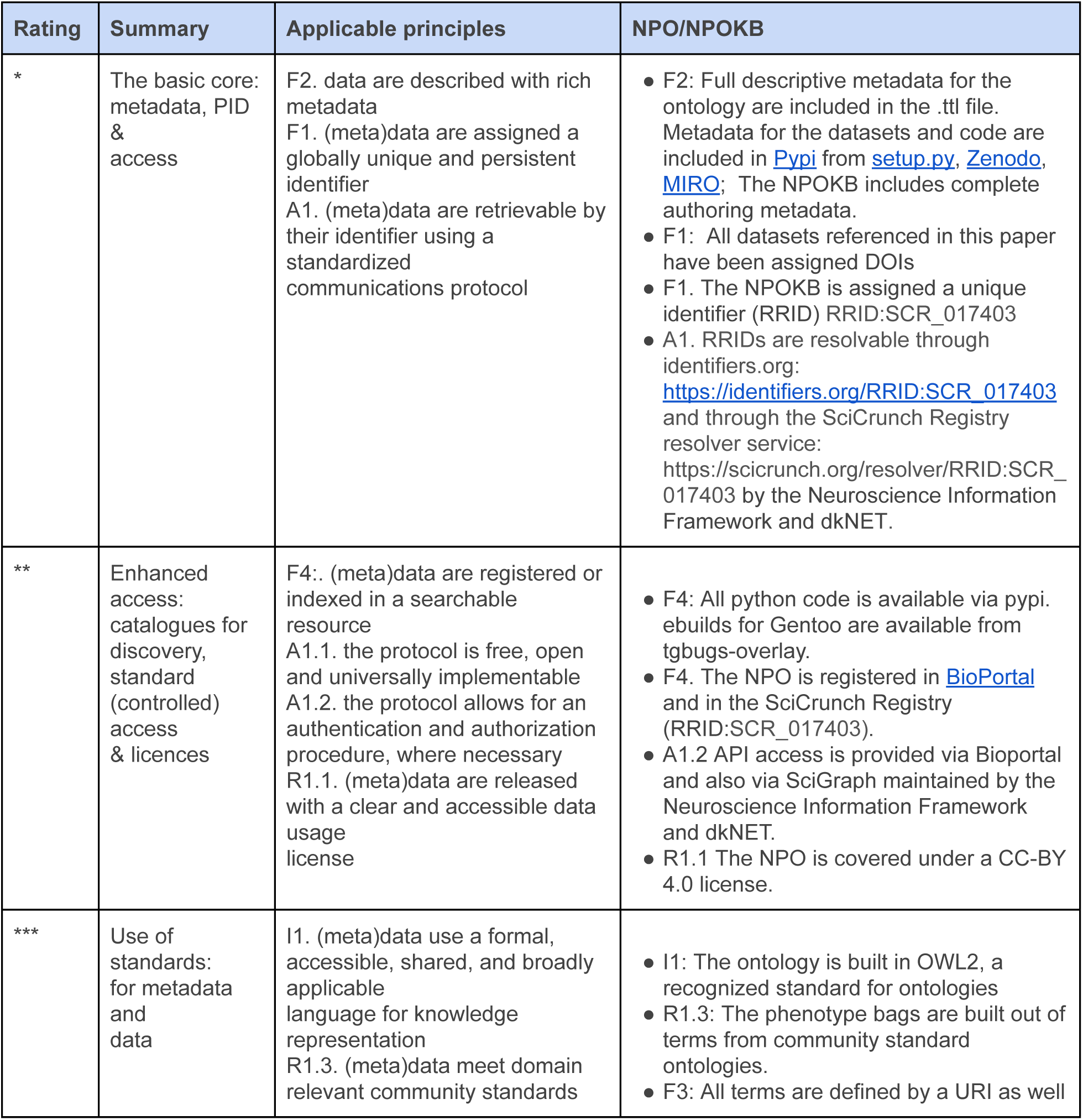
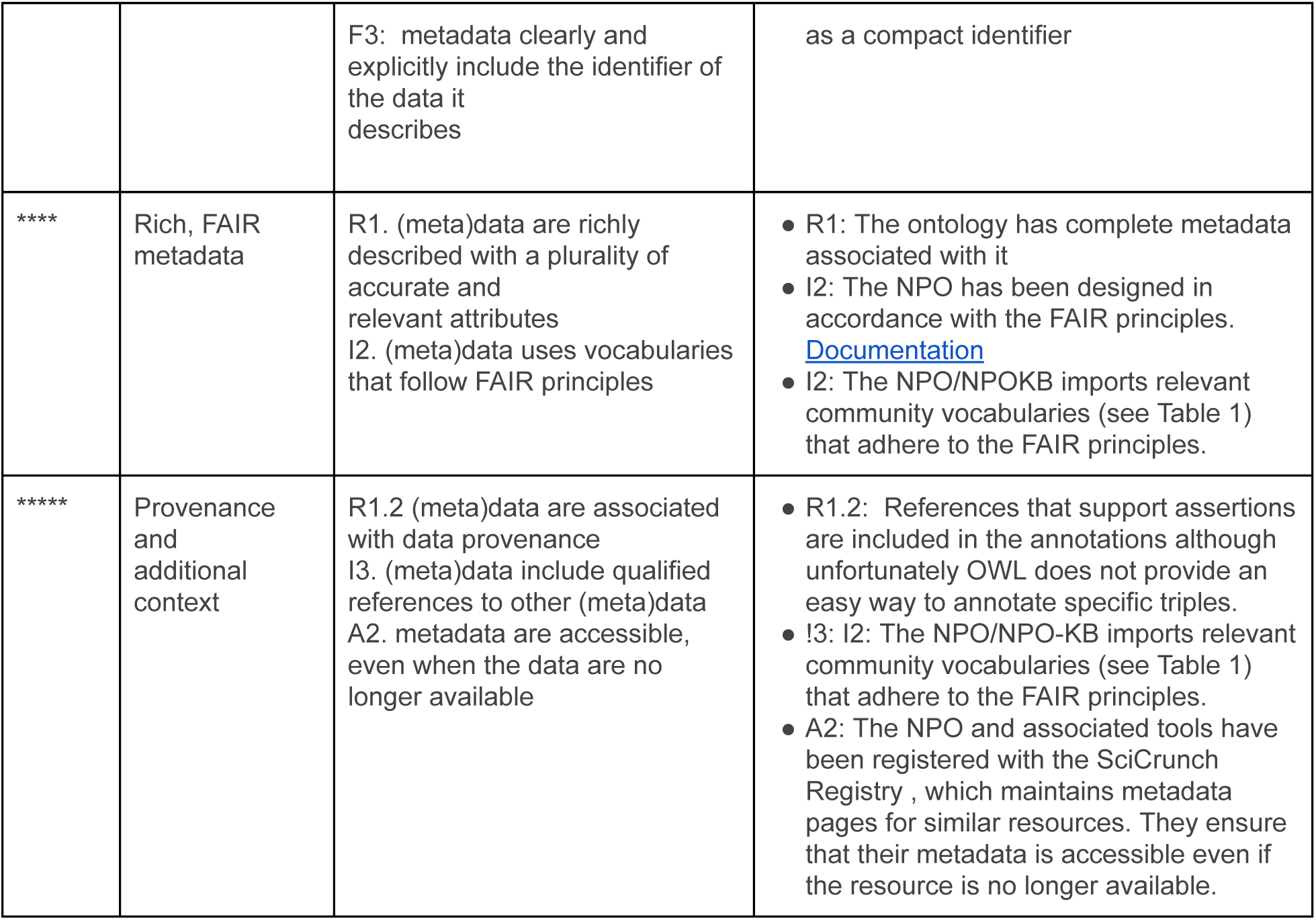
This rubric (Hodson et al., 2018) organizes the 15 FAIR principles (Applicable principles) into a hierarchical table according to how easy they are to achieve, starting from a basic core (Summary) and rates data according to level of compliance, from 1 to 4 * (Rating). We provide an evaluation of the NPO/NPOKB against these principles in column 4.

## Discussion and conclusion

The NPO provides a semantically-enriched, FAIR data model for representing the complex cellular phenotypes being generated by neuroscientists involved in individual and large scale brain initiatives. It allows the creation of machine generated taxonomies, and provides a consistent naming convention that is machine configurable. Using the NPO, we showed that we could take cellular data arising from high throughput activities, e.g., the Allen Cell Atlas, large projects like the Blue Brain Project and from individual investigators to cross between different techniques to show areas of agreement and non-alignment. This exercise is not trivial, as the multiplicity of techniques, the incomplete sampling, and the complex nomenclature present challenges. However, the NPO helps to mitigate these by allowing translation of custom lab nomenclature and experimental results into a common, semantic, and computable representation using community ontologies. The names themselves can be customized to conform to any nomenclature standard that might emerge for human consumption (e.g., Shepherd et al., 2019), but this process is managed as a formal specification rather than through agreed upon naming conventions.

We have focused our efforts on addressing the problem of cell classification vs the issue of determining neuronal types by providing a means to compare our current knowledge about cell types (our common usage types) with the many different classifications being generated by data driven methods and other experimental techniques. The distinction between a neuron type vs a neuron class is not entirely clear, and the terms are often used interchangeably. We use class here to refer to a set of neurons that satisfy a set of criteria, e.g., GABAergic neurons = all neurons that use GABA as a neurotransmitter. The number of potential classes given the number of phenotypic dimensions measured is therefore very large. Types, however, refer to neurons that are sufficiently distinct that the presence of a given set of features will reliably predict the presence of additional features that have not been measured. For example, when a cerebellar Purkinje cell is identified by a Nissl stain based on its size, shape, and location, we can reliably infer that it contains parvalbumin, and calbindin, has dendrites densely covered in dendritic spines, and uses GABA as a neurotransmitter whether or not we explicitly measure them. This definition is similar to that proposed by Zeng and Sanes (Zeng and Sanes 2017) who propose that types represent discrete groups which notionally serve a specific function while classes represent aggregates of types that share common features. Types are also the categories of cells that must be accounted for when building circuit diagrams of the nervous system (Luo, Callaway, and Svoboda 2008).

The NPO allows us to communicate about and compare measured neuronal phenotypes in a way that reflects human understanding but that can also be fully managed using modern computational methods. Genomics benefitted enormously from a community ontology for annotation of experimental results that allowed them to be communicated in a consistent and machine-processable manner, the issue of neuron typology will also benefit from a consistent annotation framework. Although there are challenges, the phenotypes themselves lend themselves to a consistent annotation framework, e.g. genes, morphological features. However, the issue of cell type itself is more fluid. Thus the NPO implements a model that distinguishes between observations in single cells (instances), proposals about cell types derived from computational analyses (EBTs) and cell types that have been recognized by one or more criteria across multiple labs and techniques (CUTs). None of these categorizations represent ground truth. Nevertheless, transcriptomics combined with data driven approaches have shown promise as a unifying technique that may allow stable cell populations to be described within a probabilistic framework (Yuste et al., 2020). Such abstractions will still likely reference entities such as brain regions, marker genes, morphology and connections and many of these will map onto well known cell types (Yuste et al., 2020). Disagreements are also still likely to arise about the nature of of these populations, particularly at finer levels of granularity NPO and the associated knowledge environment provide a bridge between such classifications generated using high throughput and integrative techniques with our accumulated knowledge over the past 100 years on cell types in the nervous system.

The work reported here should be considered a proof-of-concept; in order for the NPO to be used at the scale we envision significant additional tooling would be required. Currently, the python codes can be used now by a researcher to translate their phenotypes into NPO and they can compare their neurons locally to the NPOKB using Protégé. But to gain traction, increase ease of use and populate the knowledge base, we envision a set of on-line tools that would assist researchers in translating their phenotypes into the NPO, along with a web-accessible growing knowledge base with visualization and analysis tools for researchers to compare their neurons to what is known. Yuste and colleagues (2020) also envision an online community knowledge base where information on cell types is accumulated and linked. In addition, the NPO currently only provides the skeleton of discrete types on top of which the continuous nature of measurements needs to be integrated. Nonetheless, the goals of the BRAIN initiative and other large scale data projects are to transform our understanding of the brain through new technologies and data science and understanding the “ parts list” of the nervous system is a key objective (Zeng and Sanes, 2018). If we accept the premise that no single project or group can do it alone, then neuroscientists must produce data and knowledge artifacts like atlases and taxonomies in a way that is amenable to computation. The FAIR data principles outline some of the basic ways to do that (Table 7). Integral to FAIR is the use of community standards that make the process of searching, aggregating, and reusing data more tractable. The proposed methods do not require that we all think alike, rather, they ensure that we can employ computational methods to compare and contrast across different classification schemes. Although the proposed approaches would require a significant investment by funders and researchers alike to develop and adopt these methods, we have to measure this against the time we currently spend trying to reconcile computationally opaque and un-FAIR neuroscience data. In an ideal world, we would focus our resources on grappling with the innate complexity of the issue of cell types in the brain, rather than having to focus on reconciling the myriad number of ways we can refer to common entities in neuroscience.

## Acknowledgements

This work was supported by NIH Brain Initiative award 5U24MH114827-04, a Canadian Institute for Health Research post-doctoral fellowship and National Institutes of Health grant MH111099, the Krembil Foundation, and by funding to the Blue Brain Project, a research center of the École polytechnique fédérale de Lausanne (EPFL), from the Swiss government’s ETH Board of the Swiss Federal Institutes of Technology.

Meetings supporting this work were facilitated by the International Neuroinformatics Coordinating Facility (INCF) through the Neuroinformatics for cell types special interest group. The authors thank Felix Schürmann and Karin Holm from the Blue Brain Project for helpful comments.

## Footnotes

1 https://www.ncbi.nlm.nih.gov/taxonomy

2 https://github.com/SciCrunch/NIF-Ontology/blob/master/docs/brain-regions.org

3 https://github.com/SciCrunch/NIF-Ontology

4 https://www.ncbi.nlm.nih.gov/gene

5 https://www.ebi.ac.uk/chebi/

6 https://proconsortium.org/

7 http://geneontology.org/

8 https://github.com/tgbugs/pyontutils/blob/cc538d9c790d607cbc8c2af8a3c25f1bfa3bfc0b/ttlser/docs/ttlser.md

9 https://bioportal.bioontology.org/

