## Supplemental Materials for "The Neuron Phenotype Ontology: A FAIR Approach to Proposing and Classifying Neuronal Types"

### Table S1: Criteria used to identify a CUT in the literature

1. The cell type is commonly referenced in introductory textbooks and teaching materials for neuroanatomy, cellular and systems neuroscience and neurophysiology.
2. The cell type is referenced in multiple articles that contain experimental findings from more than one modality and more than one laboratory or research project.
3. The cell type is generally recognized in multiple mammalian species.
4. The cell type is annotated and referenced in on-line knowledge bases, e.g., NeuroElectro.org[10], Hippocampome.org[11]
5. The cell type is referenced in on-line databases of experimental data, e.g., Blue Brain Project[12], NeuroMorpho[13].

### Table S2: Abbreviations used in this paper. For transgenes and brain regions we provide links to additional information on their meaning where possible.

| Abbreviation | Full name |
|---|---|
| Adcy1 | Gene: adenylate cyclase 1 |
| Adcy2 | Gene: adenylate cyclase 2 |
| Adcy8 | Gene: adenylate cyclase 8 |
| Adcy9 | Gene: adenylate cyclase 9 |
| Adm | Gene: adrenomedullin |
| Ai14(RCL-tdT) | Trangene: Ai14 is a Cre reporter allele designed to have a loxP-flanked STOP cassette preventing transcription of a CAG promoterdriven red fluorescent protein variant (tdTomato) - all inserted into the Gt(ROSA)26Sor locus. Ai14 mice express robust tdTomato fluorescence following Cre-mediated recombination. Used in Allen Cell Types Database |
| Ai65(RCFL-tdT) | Trangene: Target a Flp and Cre-dependent tdTomato expression cassette to the mouse Rosa26 locus used in Allen Cell Types Database (https://www.addgene.org/61577/) |
| Arhgef10 | Gene: Rho guanine nucleotide exchange factor 10 |
| BICCN | Brain Initiative Cell Census Network |

[10] https://neuroelectro.org/
[11] http://hippocampome.org/php/index.php
[12] http://microcircuits.epfl.ch/#/article/article_3_mph
[13] http://neuromorpho.org/

| | |
|---|---|
| Calb | Gene:calbindin 1 |
| Calb2 | Gene:calbindin 2 (calretinin) |
| Calca | Gene:calcitonin |
| CCK, Cck | Cholecystokinin |
| Cckbr | Gene:cholecystokinin B receptor |
| CCKC | Cholecystokinin small basket cells (Paul et al., 2017) |
| Chrm2 | Gene:muscarinic acetylcholine receptor M2 |
| Chrm3 | Gene:cholinergic receptor, muscarinic 3, cardiac |
| CNP | Gene:C-type natriuretic peptide 3 |
| Cnr1 | Gene: protocadherin alpha 4 |
| Cort | Gene:cortistatin |
| Cplx2 | Gene: complexin 2 |
| Cplx3 | Gene: complexin-3 |
| CR | Calretinin |
| Crh | Gene: corticotropin-releasing factor isoform 2 precursor |
| CQ | Competency query |
| CUT | Common usage type |
| DL | Description logic |
| EBT | Evidence based type[14] |
| Edn3 | Gene: endothelin 3 |
| Esrrg | Gene: estrogen-related receptor gamma |
| GABA | Gamma amino butyric acid |
| GABAR | GABA receptor |
| Gabra1 | Gene: gamma-aminobutyric acid type A receptor subunit alpha1 |
| Gabrd | Gene: gamma-aminobutyric acid (GABA) A receptor, subunit delta |
| Gria1 | Gene: glutamate ionotropic receptor AMPA type subunit 1 |
| Gria4 | Gene: glutamate ionotropic receptor AMPA type subunit 4 |
| GluR | Glutamate receptor |
| Gpr88 | Gene:G-protein coupled receptor 88 |
| Grin3a | Gene:glutamate ionotropic receptor NMDA type subunit 3A |
| Gucy1a1 | Gene:guanylate cyclase 1 soluble subunit alpha 1 |

[14] In the data and documentation evidence based types are referred as evidence based models or EBMs which was their working name. As used here terms are equivalent.

| | |
|---|---|
| Gucy1b1 | Gene:guanylate cyclase 1 soluble subunit beta 1 |
| Hcrtr1 | Gene:hypocretin (orexin) receptor 1 |
| Hs6st3 | Gene: heparan-sulfate 6-O-sulfotransferase 3 |
| Htr2c | Gene:5-hydroxytryptamine (serotonin) receptor 2C |
| Htr3a | Gene: 5-hydroxytryptamine receptor 3A |
| Igf1 | Gene: insulin-like growth factor I |
| Kv3 | Gene: potassium voltage-gated channel, shaker-related subfamily, member 3 |
| Rspo2 | Neuropeptide. This appears as Rspn in (Paul et al., 2017; Table S7) which we suspect is a typo. |
| Zip1 | Gene:transcription factor Zip1 |
| Znt3 | Gene:zinc transporter 3 |
| Inhbb | Gene:inhibin beta-B |
| LPC | Long projection cell of Paul et al. (2017) |
| Mef2c | Gene: myocyte-specific enhancer factor 2C |
| MNC | Martinotti cells (Paul et al., 2017) |
| Nefh | Gene: neurofilament, heavy polypeptide |
| NIF | Neuroscience Information Framework |
| NIFSTD | NIF Standard Ontology (Bug et al., 2008) |
| NOS | Gene:Nitric oxide synathase |
| NOS1-creER | Transgene: Also known as nNOS-CreER-KI knock-in allele, NOS1-creER was designed to both abolish neuronal nitric oxide synthase 1 (Nos1) gene function and direct CreERT2 fusion protein expression to nNOS positive GABAergic neurons by the endogenous Nos1 promoter/enhancer elements. (https://www.jax.org/strain/014541) |
| NOS1, Nos1 | Nitric oxide synthase 1 |
| NPO | Neuron Phenotype Ontology |
| NPOKB | Neuron Phenotype Ontology Knowledge Base, also the prefix on BioPortal. |
| npokb | Compact identifier prefix for NPO identifiers. |
| NPY, Npy | Neuropeptide Y |
| Npy1r | Gene: neuropeptide Y receptor Y1 |
| Opn3 | Gene: opsin 3 |

| | |
|---|---|
| Oxtr | Gene: oxytocin receptor |
| Pde1a | calcium/calmodulin-dependent 3',5'-cyclic nucleotide phosphodiesterase 1A |
| Pde2a | Gene:phosphodiesterase 2A, cGMP-stimulated |
| Pde7b | Gene: cAMP-specific 3',5'-cyclic phosphodiesterase 7B |
| Penk | Gene: proenkephalin-A gene |
| Pnoc | Gene: prepronociceptin |
| Pparg | Gene: peroxisome proliferator-activated receptor gamma |
| Ppargc1a | Gene: PPARG coactivator 1 alpha |
| Prkg2 | Gene: protein kinase, cGMP-dependent, type II |
| Prok2 | Gene: prokineticin 2 |
| Ptn | Gene: pleiotrophin |
| PV | Parvalbumin |
| PV-cre | Parvalbumin cre line |
| Pvalb-T2A-FlpO | Transgene: Also Known As:Pvalb-T2A-FlpO-D , Pvalb-2A-FlpO-D; Pvalb-T2A-FlpO-D (Pvalb-2A-FlpO-D) mice have both endogenous gene and optimized FLP recombinase (FlpO) gene expression directed to Pvalb-expressing cells by the endogenous promoter/enhancer elements of the parvalbumin (locus.https://www.jax.org/strain/022730) |
| PVBC | Parvalbumin basket cell (Paul et al. 2017) |
| Rasl11b | Gene: RAS-like family 11 member B |
| Rgs12 | Gene: regulator of G-protein signaling 12 |
| Rgs4 | Gene: regulator of G-protein signaling 4 |
| Rgs7 | Gene:regulator of G-protein signaling 7 |
| Rln1 | Gene: prorelaxin H1 |
| RSPd2/3 | ABA Brain region: Retrosplenial area, dorsal part, layer 2/3 Structure in the Mouse Brain Atlas |
| S1 | Primary somatosensory cortex |
| S7 | Supplemental table 7 from Paul et al (2017) |
| Slc7a3 | Gene: solute carrier family 7 (cationic amino acid transporter, y+ system), member 3 |
| Slit2 | Gene: slit guidance ligand 2 |

| | |
|---|---|
| Slit3 | Gene: slit guidance ligand 3 |
| SPARC | Stimulating Peripheral Activity to Relieve Conditions |
| Sst-IRES-Cre | Transgene: somatostatin cre line developed by the Huang lab and used in Allen Cell Types Database |
| SST-flp, SST | Transgene: somatostatin-ires-Flpo (Huang neurons). Same as Sst-IRES-FlpO |
| Sst-IRES-FlpO | Transgene: FLPo recombinase expression in GABAergic neurons derived from the medial-ganglionic eminence (MGE) - as directed by the endogenous somatostatin promoter developed by Huang lab and used in the Allen Cell Types Database |
| Sst, SS | Somatostatin |
| Syt10 | Gene: synaptotagmin 10 |
| Syt4 | Gene: synaptotagmin 4 |
| Syt5 | Gene: synaptotagmin 5 (synaptotagmin 9) |
| Syt6 | Gene: synaptotagmin 6 |
| Tac1 | Gene: tachykinin precursor 1 |
| Tac2 | Gene: tachykinin 2 |
| Tacr1 | Gene: tachykinin receptor 1 |
| Trpc6 | Gene: transient receptor potential cation channel subfamily C member 6 |
| Unc5d | Gene: unc-5 netrin receptor D |
| VIP | Vasoactive intestinal polypeptide |
| Vipr1 | Gene: vasoactive intestinal peptide receptor 1 |
| Vipr2-IRES2-Cre | Transgene: Vipr2-IRES2-Cre-D knock-in mice (https://www.jax.org/strain/031332) In subcortical areas, this Cre line has expression in arcuate hypothalamic nucleus, dorsal lateral geniculate nucleus, ventral nucleus of the thalamus, bed nuclei of the stria terminalis, and scattered expression in superior colliculus. In the cortex, there is scattered expression in neurons and non-neuronal cells. (from Allen Cell Types Database documentation) |
| VISl5 | ABA Brain Region: Lateral visual area, layer 5 |
| VISp6a | ABA Brain Region: Primary visual area, layer 6a |
| VISrl2/3 | Allen Brain Atlas:Rostrolateral area, layer 2/3 |
| Wnt2 | Gene: Wnt-2 protein |

*Table S3: Comparison with other Cell Ontologies in Bioportal*

| Ontology | Description | Ontology features | NPOKB features |
|---|---|---|---|
| Cell Ontology[15] | The Cell Ontology (CL) is designed as a structured controlled vocabulary for cell types. This ontology was constructed for use by the model organism and other bioinformatics databases, where there is a need for a controlled vocabulary of cell types. This ontology is not organism specific it covers cell types from prokaryotes to mammals. However, it excludes plant cell types, which are covered by PO. See the website (From BioPortal[16]) | • Covers all cell types<br>• Built on BFO; all neurons are direct children of electrically active cell<br>• Makes use of OWL axioms<br>• Anatomical location assigned to entire cell using the "Part of" relationship<br>• Imports UBERON, CHEBI, Protein Ontology, GO<br>• Makes extensive use of GO biological processes to related molecules to cell, e.g., Neurotransmitter captured through GO biological process "neuron and ('capable of' some 'acetylcholine secretion, neurotransmission')"<br>• NPOKB CUTs should be imported into CL | • Focused on neurons<br>• Built on BFO; Neuron CUTs and EBTs are explicit classes under Neuron<br>• Makes use of OWL axioms<br>• Anatomical location assigned by location of soma using the "HasSomaLocatedIn" property<br>• Uses UBERON for anatomical locations<br>• Uses more direct relationships to relate molecules to cell, e.g., Neurotransmitter assigned by property "haNeurotransmitter" |
| Provisional Cell Ontology | Provisional cell ontology (PCL) is based on results of transcriptomic and developmental studies of brain largely carried out at the Allen Institute for Brain Science (Hodge et al. 2019) | • All cells are children of OWL: Thing; not asserted to be cells<br>• Captures developmental information<br>• No OWL axioms<br>• Does not model molecular phenotypes<br>• Uses UBERON for anatomical locations | • All neurons are children of "Neurons"> Nervous system cell > cell<br>• Makes use of OWL axioms<br>• Uses UBERON for anatomical locations + brain atlas regions<br>• Molecular phenotypes modeled through OWL axioms<br>• Does not capture developmental phenotypes |
| NeuroMorpho Brain Region Ontologies[17] | The species, brain regions and cell types are integrated into a single ontology for enabling OntoSearch functionality to mine the latest v7.0 release | • All cells are children of OWL:thing<br>• No OWL axioms<br>• Minimum number of properties<br>• Does not model molecular or | • All neurons are children of "Neurons"> Nervous system cell > cell<br>• Makes use of OWL axioms |

[15] http://cellontology.org
[16] https://bioportal.bioontology.org/ontologies/CL
[17] https://bioportal.bioontology.org/ontologies/NMOBR/?p=summary

| | | | |
|---|---|---|---|
| | of NeuroMorpho.Org. (from BioPortal) | location phenotypes<br>● Has invertebrate neurons | ● Uses UBERON for anatomical locations + brain atlas regions<br>● Molecular phenotypes modeled through OWL axioms<br>● No invertebrate neurons |

*Table S3 legend: Cell ontologies similar to NPO in Bioportal. Ontology = name of ontology; Description = description of ontology; Ontology features= Some features from the other ontologies; NPOKB features = corresponding features of NPO to highlight similarities and differences. Of the three, NPO is closely aligned with the Cell Ontology* (Diehl et al. 2016) *in design in that they are built from the BFO upper ontology and import many of the same community ontologies. They do make different modeling decisions, however, on key phenotypes, so they are not entirely interoperable. It is also similar in scope to the Provisional Cell Ontology* (Hodge et al. 2019) *in that it focuses on cell types derived from experimental techniques. However, the former does not make as extensive use of community ontologies, and very little use of the logical features of OWL to aid in classification. BFO = basic formal ontology*[18]

### Supplemental methods:

#### Working with the NPO

NeuronDM depends on a set of core ontology files that define the predicates and the different types of neuronal phenotypes that can be used to describe a neuron type. These files are packaged as part of NeuronDM releases. It is possible to use most of the NeuronDM tooling without a local copy of the NIF-Ontology repository. For development NeuronDM requires a local copy of the NIF-Ontology repository with the neurons branch[19] checked out.

Detailed documentation on how to load the NPO into the Protégé Ontology Tool[20] is maintained in GitHub.[21] However, the NIFSTD is a very large ontology with multiple import statements that can be difficult to work with. First, loading the full ontology in Protégé requires retrieving all files in the import chain and patching the ontology catalog file and any external ontologies with known issues. Once this is done, it is possible to load the ontology and run the reasoner. The full NIFSTD ontology is relatively large and reasoning over it and the full set of predicates available for describing phenotypes requires roughly 4 gigs of memory at the outset which grows as the reasoning proceeds. On a quad-core machine from 2013 loading takes about 30 seconds and initial reasoning about another 10 seconds, however incremental reasoning thereafter can take many minutes. In practice we have found that it can sometimes be more expedient to close Protégé, reload NIFSTD, and reason from scratch than to use incremental

[18] https://basic-formal-ontology.org/#:~:text=The%20Basic%20Formal%20Ontology%20(BFO,is%20a%20genuine%20upper%20ontology.&text=BFO%20is%20used%20by%20more,driven%20endeavors%20throughout%20the%20world.
[19] https://github.com/SciCrunch/NIF-Ontology/blob/master/docs/Neurons.md
[20] https://protege.stanford.edu/
[21] https://github.com/SciCrunch/NIF-Ontology/blob/master/docs/Neurons.md#protege-development-setup

reasoning when there are changes. The fully expanded and reasoned NIF ontology is also accessible via a SciGraph API endpoint (see using NIFSTD for details) that serves a graph built using an automated version of the loading process. SciGraph is a Neo4j backed ontology store that is used by the python representation of the NPO to obtain lexical information about phenotypes and to verify that they are in the set of known terms if the term is not part of the core classes that are stored locally.

To make the NPO more accessible for inspection and use by subject matter experts, we created NPO "Light", a stripped down .ttl version that loads in most desktop installations of the Protégé Ontology Browser (version 5.5 or higher)[22] without the requirement to download the full NIFSTD or any additional modifications to the local machine. NPO Light can be loaded into Protégé on either a PC or Mac using the "Open from URL" option in the file menu. NPO Light  supports basic reasoning about neuron types, but does not support the full reasoning chain.  In addition, the NPO Light file includes the genes, proteins and brain regions that are used in the current set of neuron phenotype definitions, i.e., NPO Light does not import the full gene list of NCBI gene. Unless otherwise specified, all of the examples presented in this work were derived from NPO Light.

### Modeling Location Phenotypes

For location based queries the objective is to be able to find neurons that are located in some region. Human beings automatically fill in the implied "or any part of that region," however, the ontology cannot do this automatically. To accomplish we do two things. First we lift phenotypes of the that are defined using location predicates of the form =?subject hasLocationPhenotype: :Location= into the form =( hasLocationPhenotype: SOME ( partOf: SOME :Location ) )=. Second we add a partOfSelf axiom =:Location SubClassOf: ( partOf: SOME :Location )= to all anatomical entities that are used as phenotype values. This solution provides query behavior that matches what human beings expect when they write a DL query of the form =hasPhenotype some ('part of' some Location)=. This structure is also compatible with both the FaCT++ and ELK reasoners. A query of the form =hasSomaLocatedIn some Location= does work if a subPropertyChain axiom of the form =hasLocationPhenotype: owl:propertyChainAxiom (hasLocationPhenotype: partOf: )= is added to the ontology, however this only works with the ELK reasoner since inclusion of that axiom crashes FaCT++. For more details on the modelling of location phenotypes please see the documentation on OWL modelling decisions [23].

#### Atlas regions vs anatomical names

A continuing challenge in building ontologies for neuroscience has been the large number of anatomical terminologies derived from brain atlases and parcellation schemes that must be reconciled when referring to anatomical locations. The anatomical locations for CUTs are expressed in generic anatomical terms using the pan-species UBERON ontology (Mungall et al. 2012). Because EBTs and instances are tied directly to experimental data, locations may be expressed in terms of a particular brain atlas rather than a generic anatomical term, e.g., the

---

[22] https://raw.githubusercontent.com/SciCrunch/NIF-Ontology/neurons/ttl/neuron-development.ttl.

[23] https://github.com/tgbugs/pyontutils/blob/master/neurondm/docs/basic-model.org

Paxinos and Franklin mouse brain atlas (Franklin and Paxinos 2008) or the Allen Mouse Brain Reference Atlas (RRID:SCR_013286) terminology.

For the NPO, we enhanced NIF's core anatomical ontologies by creating a parcellation scheme ontology[24]. The current version contains ~ 30 nomenclatures from major brain atlases in rat, mouse, human and cat. For each parcellation scheme, we either imported the hierarchy if it was available, e.g., Allen Brain Mouse Atlas, or created a hierarchy based on region names and information provided with the nomenclature. This process sometimes necessitated creating additional superclasses that were not present in the original nomenclature. For example, the Paxinos atlases divides the cerebellum up into subregions, but does not contain the root:cerebellum. To relate atlas structures to common brain regions, we utilize the "delineated by" relationship, e.g.,

> *Neocortex +SOM neuron = neuron and hasPhenotype some ('part of' some (neocortex or (delineates some neocortex))) and hasPhenotype some (Sst or SST or Sst-IRES-Cre or Sst-IRES-FlpO)*

This DL query returns neurons that have their soma located in any part of the neocortex or in any atlas structure that delineates a subregion of the neocortex.

[24] http://ontology.neuinfo.org/NIF/ttl/generated/parcellation-artifacts.ttl
